# A G protein coupled receptor like protein systematically modulates nutrient-growth by Ca^2+^ mediated phosphatidylcholine perception during evolutionary adaptation

**DOI:** 10.1101/2022.06.17.496194

**Authors:** Zhixiong Huang, Xue He, Xueqiang Zhao, Wan Teng, Mengyun Hu, Yijing Zhang, Junming Li, Hui Li, Yiping Tong

**Author notes:** Corresponding author. (Y.T.).

## Abstract

Nutrients are important for growth in both plants and animals, uncovering of signaling pathway in nutrients determined growth is essential. Here we cloned *TaPCGR1-3B* (*Phospholipid Coordinated Growth and nutrients Response* 1), controlled by SNPs on alternative splicing and transcription factors binding, conferring nitrogen deficiency response. TaPCGR1-3B was localized in plasma membrane and endoplasmic reticulum of meristem cell. Nitrogen deprivation stimulated interaction of TaPCGR1-3B with G protein alpha subunit and phospholipase C 9, which was inhibited by phosphatidylcholine, to trigger Ca^2+^ signaling and inhibit normal growth. Knockdown of *TaPCGR1* rescued the growth inhibition caused by nutrient deficient conditions by modulation of phosphatidylcholine induced growth gene expression through Ca^2+^ signaling disruption. Modulating of phosphatidylcholine mediated *TaPCGR1* activity thus tightly regulated growth through Ca^2+^ signaling.

---

Overuse of fertilizer since green revolution led to soil hardness and greenhouse gas (N_2_O, which is 1000 more harmful than CO_2_) release (*1, 2*), which have been deteriorating environment and finally threatening to human survival by reducing grain yield, extreme weather, perturbing water cycling, etc. (*3*). Furthermore, the grain-yield increase range per nitrogen fertilizer input after green revolution have been diminishing (*4*). However, the ever-increasing population needs more food. To address the challenge, we need to understand the mechanisms by which nitrogen deficiency affects plant growth in the coordination between genome and stresses adaptation. Although nitrate was supposed to trigger downstream genes expression by regulating Ca^2+^-sensor protein kinases (CPKs) (*5*), the nitrogen regulation network was much more complex than we thought (*6*). Some peptides were discovered to sense heterogenous nitrogen distribution in soil (*7*). However, in modern agriculture, fertilizers are evenly spread. Although CHL1/NRT1.1 sensed transient nitrate level difference (*8*) and phytohorme crosstalked with nitrogen signalings(*9-11*), signal transduction pathway involved in long time nitrogen deficiency (under agriculture trials rather than experiment condition) is unknown.

Nitrogen deficiency decreased tiller number compared with sufficient supply of nitrogen fertilizer (Fig. 1) in common wheat (*12*). We reasoned that the trade-off between nitrogen supply and tiller number is regulated by a particular molecule through unknown signaling pathway. To isolate genes that play crucial roles in long term nitrogen deficient adaptation, we construct a BC_5_F_6-7_ population derived from backcross hybrid between Xiaoyan 54 (XY54) and Jing411 (J411), which showed contrasted nitrogen deficiency adaptation (*12*). We focused on the grain yield difference in the population since grain yield (Fig. 1 A and B) and spike number (Fig. 1C), most likely to be affected under nitrogen deficient condition, rather than grain number (per spike) (Fig. 1D) and thousand grain weight (Fig. 1E) between haplotype I and haplotype II. Indeed, the tiller elongation in haplotype II was inhibited compared with haplotype I, though the tiller initiation was not affected in both haplotypes (Fig. 1F and G, S1). Bulk RNA sequencing association results from highest and lowest number of spike number indicated a genetic region near centromere in chromosome 3B was significantly associated with spike number difference (Fig. S2). We isolated a pair of near-isogenic lines (NILs) from this backcross population that was heterogenous in the region. We further narrowed down the genetic region in the heterogenous offsprings and next sequenced genomic sequences in the region between near-isogenic lines. We found an unknown gene *TaPCGR1*-3B (*Phospholipid Coordinated Growth and environment Response 1*) had eight single-nucleotide polymorphism (SNP) sites in different haplotypes (Fig. S3). One SNP in 5`-UTR, two SNPs in intron, and five SNPs in 3`region of gene body (Fig. S3).

**Fig.1.**
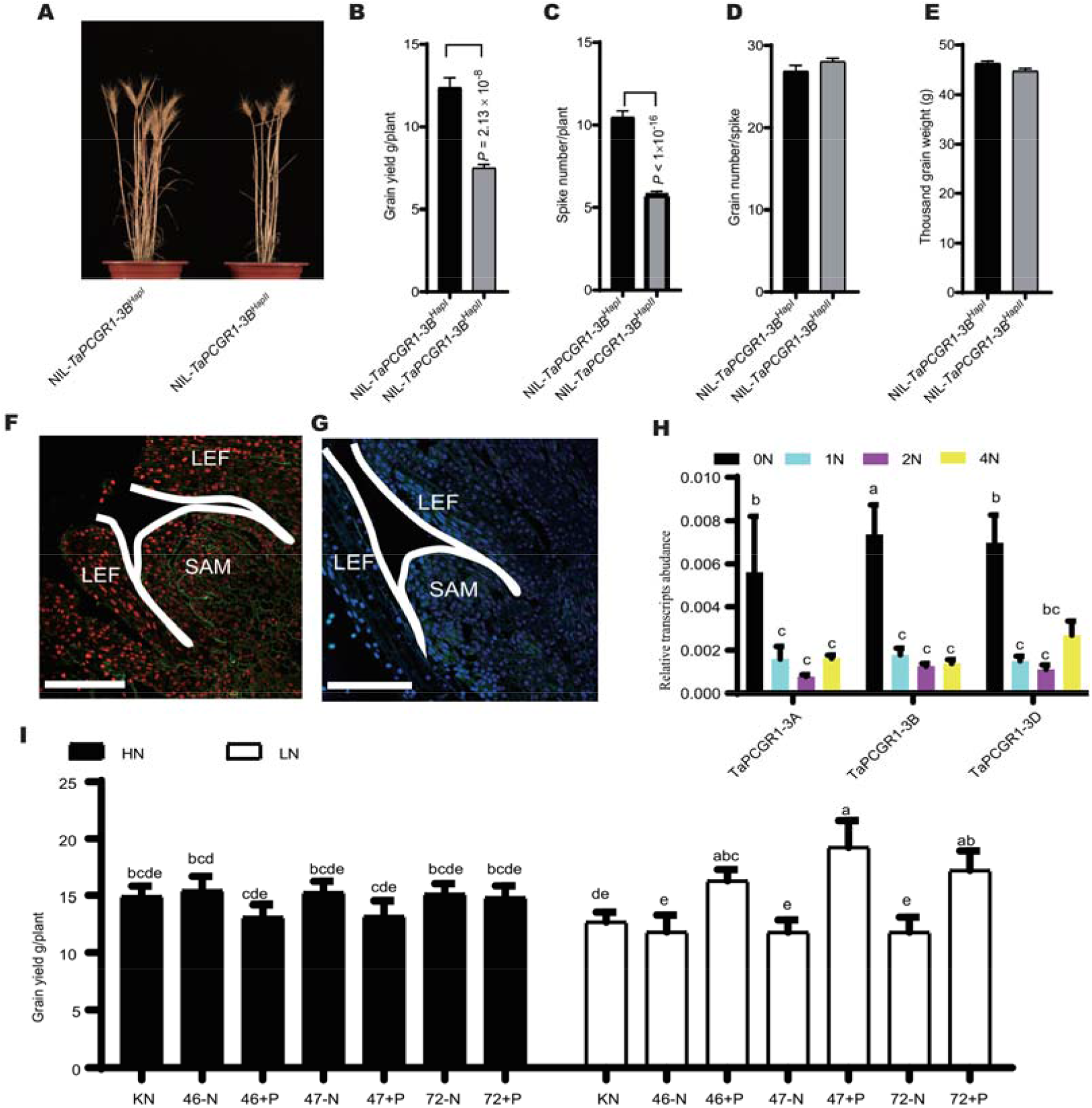
*TaPCGR1-3B* conferred nitrogen deficiency response. (**A**) Mature NIL-*TaPCGR1-3B*^HapI^ and NIL-*TaPCGR1-3B*^HapII^ grown under nitrogen deficient condition (30 kg N/ha) selected from more than four independent replicates. Isogenic lines NIL-*TaPCGR1-3B*^HapI^ and NIL-*TaPCGR1-3B*^HapII^ are generated by introducing NIL-*TaPCGR1-3B*^HapII^ from J411 into XY54 (BC5F6). Scale bar, 10 cm. (**B**) Grain yield per plant. (**C**) Spike number per plant. (**D**) Grain number per spike. **(E)** 1000-grain weight. Data in (B) to (E) are mean ± S. E. (n > 40) and *P* values from student’s *t*-test. **(F-G)** Tissue localization of TaPCGR1-3B. shoot basal parts tissue immunostaining with anti-TaPCGR1 antibody in NIL-*TaPCGR1-3B*^HapI^ **(F)** and NIL-*TaPCGR1-3B*^HapII^ **(G)**. Fluorescence from secondary antibody (green) and nucleus stained with DAPI (red and blue) are shown. Scale bars, 200 μm. Images are representative of more than 4 independent replicates. LE, leaves. SAM, shoot apical meristem. **(H)** The response of *TaPCGR1-3A, -3B* and *-3D* to nitrogen availability. The plants of KN199WT (KN199 wild type) were grown for 4 weeks at 0.0 mM NH_4_NO_3_ (0N), 0.5 mM NH_4_NO_3_ (1N) and 1.0 mM NH_4_NO_3_ (2N). The shoots were collected for gene expression analysis. The genes expression level was normalized to that of *Tatublin* as internal control. (**I**) Grain yield of KN and RNAi positive and negative lines under sufficient (HN, 270 kg N/ha) and deficient (LN,90 kg N/ha) nitrogen supply condition. Data are mean ± S. E. (n = 8). Different letters above the column indicate significant difference at *P* < 0.05 level.

*TaPCGR1*-3B is a gene closely related to *Glycerophosphocholine acyltransferase* (*GPCAT*) in yeast (*13*) (Fig. S4). We next investigated whether *PCGR1* responded to different nitrogen supply level. Indeed, *PCGR1-3A, PCGR1-3B* and *PCGR1-3D* were significantly induced by after two-weeks nitrogen deprivation treatment (Fig. 1H). To investigated the function of *PCGR1*, we created three independent RNA interference (RNAi) lines (Fig. S5). We further explored the RNAi traits under different nitrogen supply level. No significant difference in grain yield among kenong199 (KN199WT), negative control lines (46-N, 47-N and 72-N) and positive RNAi lines (46+P, 47+P, 72+P) was observed under high nitrogen supply condition. Nevertheless, positive RNAi lines had more grain yield than negative lines and KN199WT under nitrogen deficient condition (Fig. 1I). These results indicated that *PCGR1* was important in nitrogen adaptation and modulation of grain yield under long timer nitrogen deficiency conditions rather than under high nitrogen supply.

We next investigated whether the SNP in 5`-UTR affected gene expression. Allelic variation at the 5′-UTR position in the *TaPCGR1-3B* promoter region occurred at the binding site of a *bZIP* gene family transcription factor T/C-box (GACGTT). In the *in vitro* EMSA assay, the allelic variant *TaPCGR1-3B*^HapI^ corresponded to a free probe that fully competed with the biotin-bearing probe at low concentration (Fig. 2A). In contrast, the free probe corresponding to the allelic variant *TaPCGR1-3B*^HapII^ only competed for biotin-bearing with the free probes at high concentration (Fig. 2A). *In vivo* ChIP and *in vitro* EMSA assays showed that the mutant T/C-box (GAGGTT) in *TaPCGR1-3B*^HapII^ significantly reduced the binding capacity to TabZIP45-4B (Fig. 2A and B). Indeed, knockout of one of the bZIP gene family member *TabZIP45-4B*, the gene expression level of *TaPCGR1-3A* and *TaPCGR1-3B* were significantly reduced (Fig. 2C). In contrast, overexpression of *TabZIP45* significantly promoted *TaPCGR1-3B* transcription (Fig. 2D). We next accessed whether the gene expression level of *TaPCGR1-3B* under various nitrogen supply was changed. As expected, the significant difference of mRNA abundance of *TaPCGR1-3B* between NIL-*TaPCGR1-3B*^HapI^ and NIL-*TaPCGR1-3B*^HapI^ under nitrogen deprived condition was observed, whereas no significant differences between the two haplotypes were observed at other nitrogen supply level (Fig. 2E). These results indicated that SNP in 5′-UTR region affected the *TaPCGR1-3B* transcription under nitrogen deprived condition.

**Fig.2.**
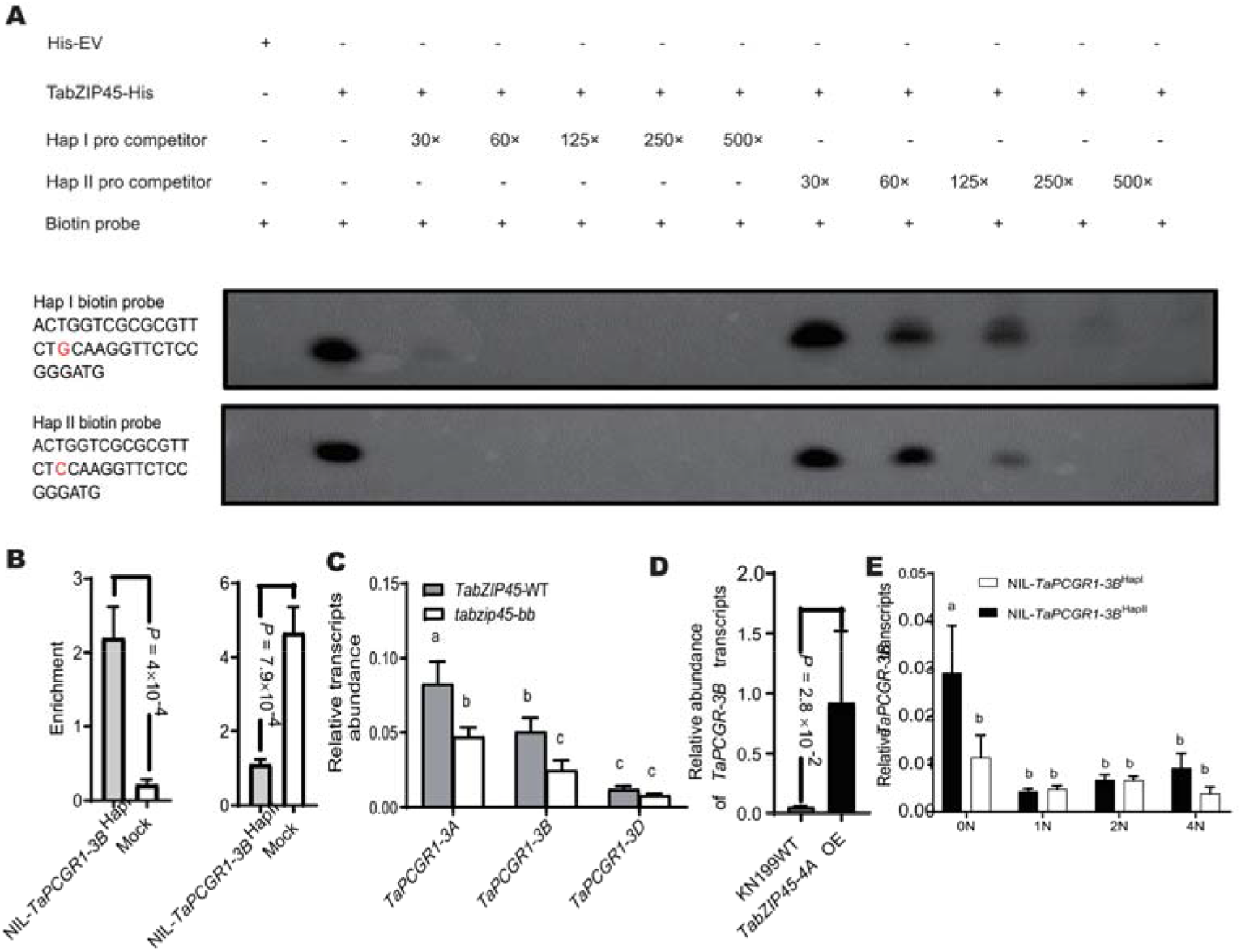
*TabZIP45* regulated *TaPCGR1-3B* transcription by binding to UTR region. (**A**) Electrophoretic mobility shift assay (EMSA) detects binding of TabZIP45 on promoter of NIL-*TaPCGR1-3B*^HapI^ and NIL-*TaPCGR1-3B*^HapII^. His-EV, Protein purified in empty vectors. TabZIP45-His, protein TabZIP45 purified with his tags. Hap I pro competitor indicates NIL-*TaPCGR1-3B*^HapI^ pro ompetitor. Hap II pro competitor indicates NIL-*TaPCGR1-3B*^HapII^ pro competitor. competitive promoter DNA fragments concentration increases from 0, 10/64, 10/16, 10/4, 10 pmol/L. Biotin (biotin probe, 20 fmol) probes promoter DNA fragments of *TaPCGR1-3B*^HapI^ (upper panel, Hap I biotin probe) and NIL-*TaPCGR1-3B*^HapII^ (down panel, Hap II biotin probe). (**B**) ChIP assays were performed with or without (MOCK) TabZIP45 antibody. NIL-*TaPCGR1-3B*^HapI^ and NIL-*TaPCGR1-3B*^HapII^ indicate fragments of the NIL-*TaPCGR1-3B*^HapI^ and NIL-*TaPCGR1-3B*^HapII^ promoters containing the SNP. Data in B are mean ± S. E. (n = 5). (**C**-**D**) Relative abundance of *TaPCGR1* transcripts. (**C**), above ground samples were collected in Hebei province under LN (45 kg N/ha) N supply at tillering stage. (**D**), Leaves were collected in Hebei province under HN (270 kg N/ha) N supply at jointing stage. *TabZIP45*-WT, homozygous wild type offspring. *tabzip45-bb*, homozygous mutant offspring. KN199WT, KN199 wild type bread wheat. *TabZIP45-4A*OE, *TabZIP45-4A* overexpression transgenics lines. Data in C are mean ± S. E. (n = 10). Data in D are mean ± S. E. (n = 4). (**E)** Relative abundance of *TaPCGR1-3B* transcripts in roots. Seedlings were grown at increasing N supply (0.00 mM NH_4_NO_3_, 0N; 0.50 mM NH_4_NO_3_, 1N; 1.0 mM NH_4_NO_3_, 2N; 2.0 mM NH_4_NO_3_, 4N). Isogenic lines NIL-*TaPCGR1-3B*^HapI^ and NIL-*TaPCGR1-3B*^HapII^ are generated by introducing *TaPCGR1-3B*^HapII^ from J411 into XY54 (BC5F6). *P* values are calculated by two tailed unpaired *t* test. Data in E are mean ± S. E. (n ≥ 6) and different letters denote statistically significant differences (*P* < 0.05) by two-way ANOVA multiple comparisons. All data are from at least three independent experiments.

Under nitrogen deficient condition, NIL-*TaPCGR1*^hapI^ had more spike number and higher survival rate than NIL-*TaPCGR1*^hapII^ (Fig. S3B and C). We next determined whether the SNPs (in introns and 3`end of gene body) affected the gene function in NIL-*TaPCGR1*^hapI^ and NIL-*TaPCGR1*^hapII^ (Fig. S3A). We isolated mRNA from NIL-*TaPCGR1*^hapI^ and NIL-*TaPCGR1*^hapII^ under nitrogen deprived conditions. Reverse Transcriptase Polymerase Chain Reaction (RT-PCR) showed that NIL-*TaPCGR1*^hapI^ had only one type of transcript, whereas NIL-*TaPCGR1*^hapII^ had at least two types of transcripts (Fig. S3D). Indeed, sequencing results indicated that the short variant and long variant encoded two different isoforms of PCGR1 (Fig. S6A). Online prediction indicated seven transmembrane domains of PCGR1 (Fig. S6 B-E). The short isoform was lack of three transmembrane domains which contained a conserved FXXXXXA/VYXL motif, which are predicted to be critical in compound binding (Fig. S6A and F). The different splicing binding factors sites in NIL-*TaPCGR1*^hapI^ and NIL -*TaPCGR1*^hapII^ produced the different splicing variants of *TaPGGR1-3B* (Fig. S3A and D). Thus, these results indicated that SNPs in intron and 3`end of gene body led to different mRNA variants of *PGGR1* by alternative splicing.

To further characterize the function of *TaPCGR1-3B*, the localization of the protein at the subcellular level was first explored. Immunoblotting experiments revealed a similar distribution pattern of *TaPCGR1* to that of the membrane protein hydrogen ion H^+^-ATPase (Fig. 3A). TaPCGR1-3B-GFP in common wheat protoplast showed a membrane structure (Fig. 3B). Indeed, subcellular localization experiments in protoplasts revealed an overlap of the green TaPCGR1-3B-GFP (green fluorescent protein) and the red PLASMA MEMBRANE INTRINSIC PROTEIN 2 (PIP2)-mCherry (plasma and endoplasmic reticulum membrane protein marker) and mCherry-HDEL (endoplasmic reticulum membrane protein marker) (Fig. 3C and D). These results indicated that TaPCGR1-3B localized in plasma and endoplasmic reticulum membrane. In addition, TaPCGR1-3B mainly expressed in shoot apical meristem (SAM) and leaves (Fig. 1F and G).

**Fig.3.**
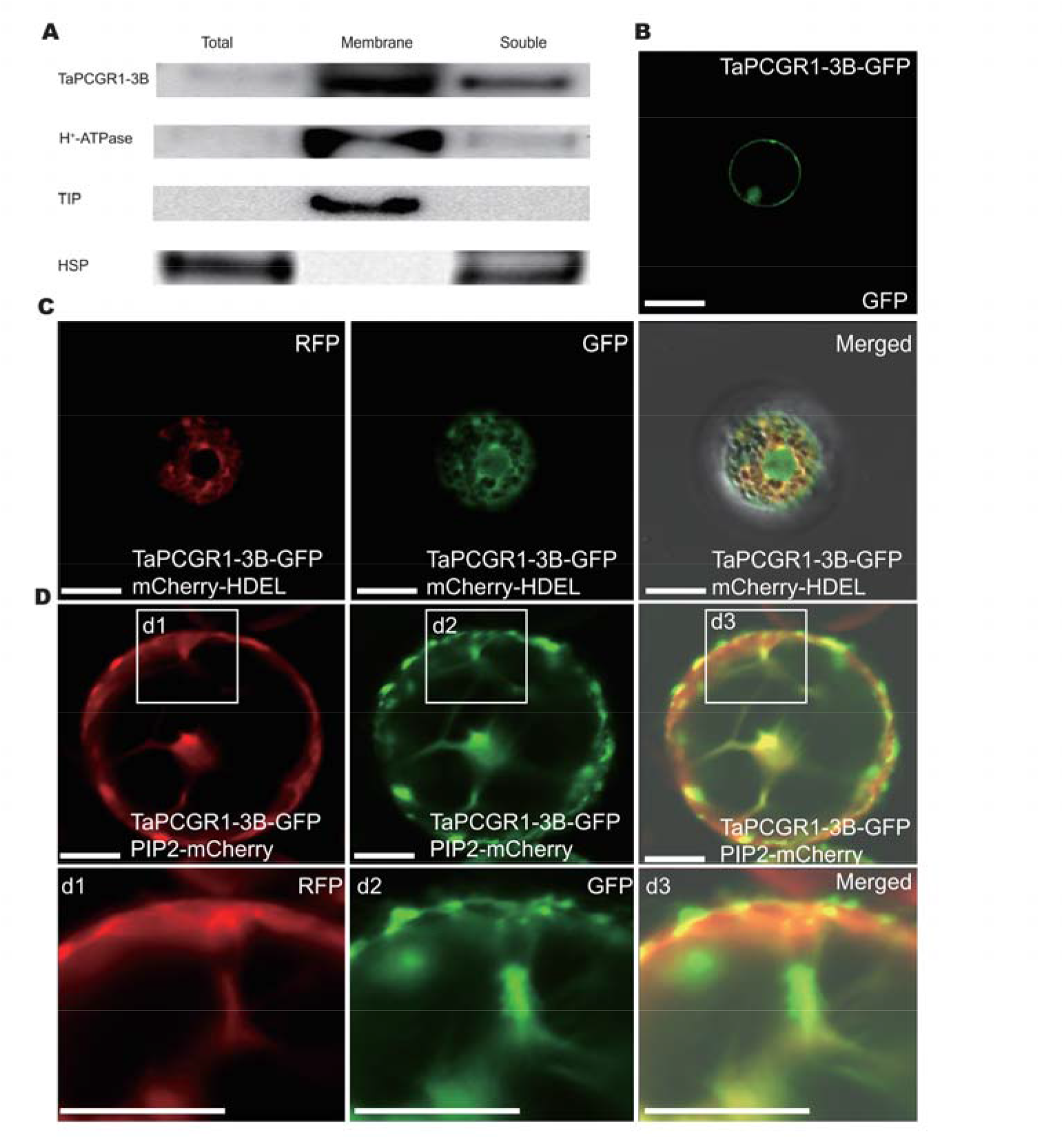
TaPCGR1-3B was localized in endoplasmic reticulum and cytoplasmic membrane. **(A)** Immunoblotting of PCGR1, H^+^-ATPase (plasma membrane marker), TIP (Tonoplast marker) and HSP (soluble protein marker) *in vivo*. **(B)** Membrane localization of TaPCGR1-3B in wheat protoplast cells. **(C)** Co-localization of TaPCGR1-3B-GFP (green) and endoplasmic reticulum (ER) marker (HDEL-mCherry) in wheat protoplast cells. **(D)** Co-localization of TaPCGR1-3B-GFP (green) and cytoplasmic and ER membrane marker plasmid (PIP2-mCherry) in tobacco (*Nicotiana tabacum*) cell. The d1, d2 and d3 indicate the enlargement of parts inside the white rectangle above respectively. Scale bars, 50 μm.

To further uncover the function of *TaPCGR1*, we found that TaPCGR1 was predicted to be very closely related to G protein coupled receptors (GPCR) (Fig. S4 and S6). We screen the protein interaction database and found that phospholipase C 9 (PLC9) interacted with GPCAT (*14*). To date, we know of only one G protein alpha subunit (GPA) in the plant kingdom. Therefore, we next tested whether TaPCGR1-3B interacted with PLC9 and GPA. As expected, TaPCGR1-3B interacted with PLC9 and GPA *in vivo* and *in vitro*, respectively, but not with the premature stop-gain mutated PLC9 (Fig. 4). Unexpectedly, TaPCGR1-3B interacted with the short splice variant encoding TaPCGR1-3B (Figure S7), indicating that the protein encoded by the short splice variant still interacted with effector protein to trigger downstream events. Thus, TaPCGR1-3B interacted with PLC and GPA.

**Fig.4.**
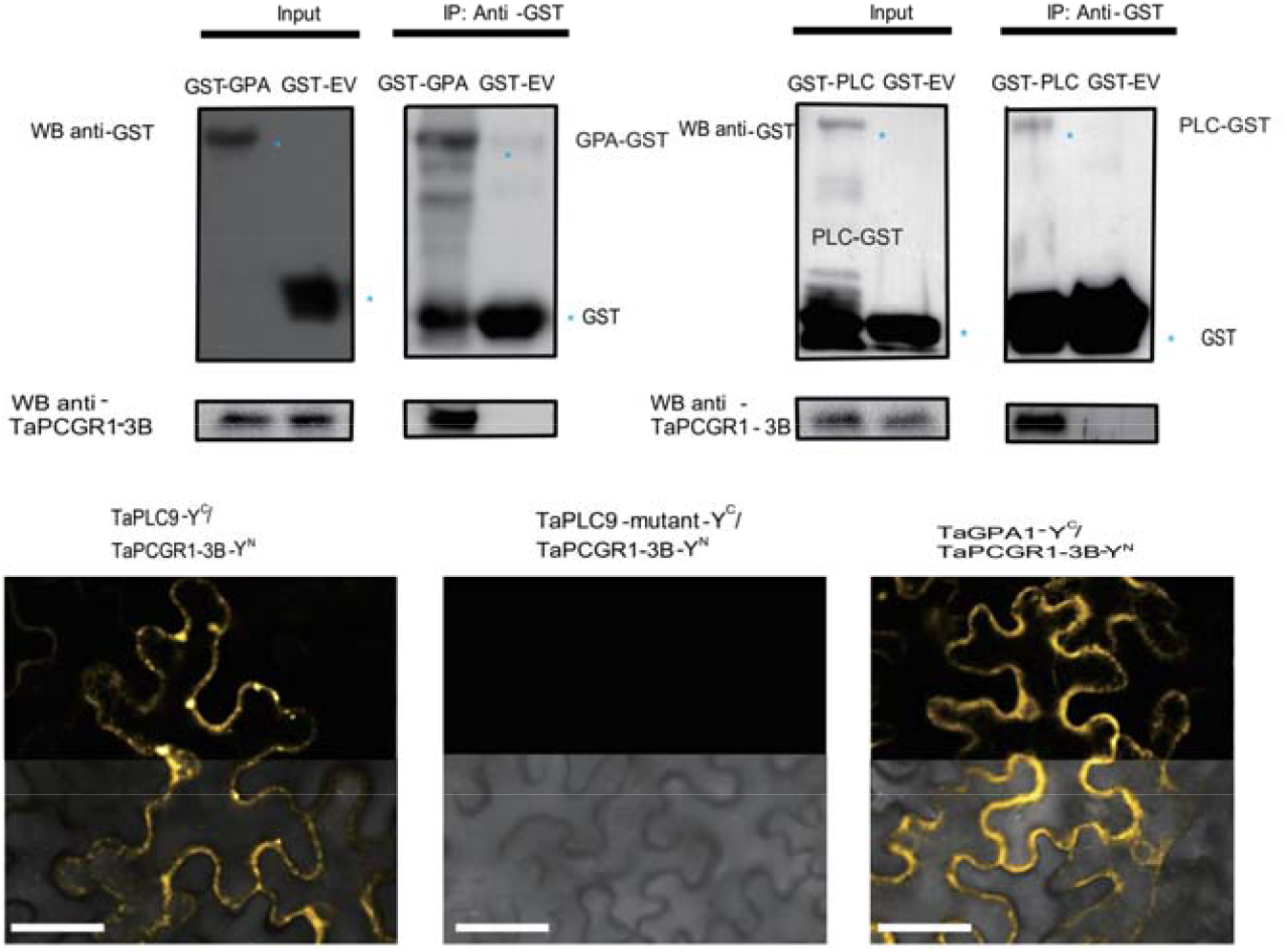
TaPCGR1-3B was targeted by TaPLC9 and TaGPA1. **(A)** Co-immunoprecipitation assay: Purified GST-GPA1 or GST-EV was incubated with KN199WT root tissue lysate, and then was detected by western-blot. **(B)** Co-immunoprecipitation assay: Purified GST-PLC or GST-EV was incubated with KN199WT root tissue lysate, and then was detected by western-blot. IP, immunoprecipitation. WB, western blot. *, GST signal. **(C-E)** BiFC experiment: gene1-YFP C terminal part (Y_C_) and targeting gene2-YFP N terminal part co-expressed in tobacco (*Nicotiana tabacum*) leaves and then was scanned with confocal microscope: (C) **Ta**PLC9-Y^C^/**Ta**PCGR1-Y^N^, (D) **Ta**PLC9-mutant (C terminal deletion mutant)-Y^C^/**Ta**PCGR1-Y^N^ and (E) **Ta**GPA1-Y^C^/**Ta**PCGR1-Y^N^. Scale bar, 100 μm.

The interaction of TaPCGR1-3B with PLC and GPA indicated that TaPCGR1-3B may function as GPCR. Indeed, online prediction by machine learning and big public database mining indicated the binding affinity between phospholipid and TaPCGR1-3B. Furthermore, previous research indicated that the GPCAT, orthologue of TaPCGR1-3B, was an enzyme involved in phosphatidylcholine (PC) synthesis-hydrolysis metabolism (*10*). We next determined whether the PC affected the interaction of TaPCGR1-3B with PLC and GPA. The addition of PC significantly reduced the interaction of TaPCGR1-3B with either PLC or GPA (Fig. S8A and B). To further addressed the function of phospholipids in downstream gene expression regulation, we characterized *Tatab1*, a *WUSCHEL* ortholog *tab1* (*TILLERS ABSENT 1*, responsible for both tiller formation and elongation) (15), transcriptional stimulation by PC, β-Acetyl-γ-O-alkyl-L-α-phosphatidylcholine (PAF), hexadecyl azelaoyl phosphatidylcholine (Azelaoyl PAF) and Phosphatidylethanolamine (PE) in NIL-*TaPCGR1-3B*^hapI^ and NIL-*TaPCGR1-3B*^hapII^, respectively (Fig. S8C and D). Differences in the transcript levels of *Tatab1* induced by the addition of different phospholipid were not significant in NIL-*TaPCGR1-3B*^hapI^, whereas only PC significantly increased the *Tatab1* transcription in NIL-*TaPCGR1-3B*^hapII^ compared with other types of phospholipids (Fig. S8C and D).

To bolster evidence that *TaPCGR1-3B* was important in lipids signaling, several lipids were tested for *Tatab1* activation. All phospholipids tested promoted tab1 expression in haplotype I compared to haplotype II (Figure S9). Furthermore, we found that *Tatab1* transcriptional level was increased in RNAi positive line under nitrogen deficient condition compared with negative line (Figure S10). These results indicated that the elevated expression of *Tatab1* in haplotype I compared with haplotype II was due to the insensitivity of *TaPCGR1-3B* in NIL-*TaPCGR1-3B*^hapI^ to PC, which inhibited the downstream events triggered by interaction between *TaPCGR1-3B* and effector proteins.

Nitrogen deficiency induced phospholipids degradation (*16, 17*). The disturbance of phospholipids was reported to trigger calcium signaling (*18, 19*). We hypothesized that under nitrogen deficiency conditions, PC hydrolysis stimulates the interaction of TaPCGR1-3B with effector proteins, which in turn promotes calcium signaling and ultimately inhibits tillering, and vice versa. Indeed, we found that the remove from nitrogen sufficient conditions (HN) to conditions without nitrogen supply (0N) significantly promoted Ca^2+^ influx, whereas removing from nitrogen deficient conditions (LN) to 0N only slightly promoted Ca^2+^ influx in NIL-*TaPCGR1-3B*^hapII^ (Fig. S11 A, C and E). These results indicated that nitrogen supply levels difference stimulated calcium signaling in NIL-*TaPCGR1-3B*^hapII^. In contrast, calcium outflux rather than influx were observed in both removing from HN to 0N and LN to 0N in NIL-*TaPCGR1-3B*^hapI^ (Fig. S11 A, C and E). Moreover, the outflux between HN to 0N and LN to 0N was not significant in NIL-*TaPCGR1-3B*^hapI^, indicating that NIL-*TaPCGR1-3B*^hapI^ was less sensitive to nitrogen supply levels difference than NIL-*TaPCGR1-3B*^hapII^ (Fig. S11 A, C and E).

To further elucidate the function of *TaPCGR1-3B* in calcium signaling, we next accessed the RNAi positive line and negative control line of *TaPCGR1-3B* in responsive to different nitrogen supply. The positive line had less Ca^2+^ influx in both HN to 0N and LN to 0N removing compared with negative line, indicating that the *TaPCGR1-3B* was important in nitrogen supply levels sensation (Fig. S11 B, D and E).

We next evaluated the Ca^2+^ signal localization in both haplotypes under different nitrogen supply conditions. Under sufficient nitrogen supply level (HN), no Ca^2+^ fluorescent signal was observed in the two haplotypes (Fig. S11 F and H). Under nitrogen deficient conditions, however, positive signal was only observed in NIL-*TaPCGR1-3B*^hapII^ but not in NIL-*TaPCGR1-3B*^hapI^ (Fig. S11 H, I and J). The calcium fluorescent signal was localized in the boundary of meristem and elongation zone, indicating the calcium was important in cell division (Fig. S11 I and J). Taken together, *TaPCGR1-3B* was important in calcium signaling, which was induced by nitrogen deficiency.

We showed that the phospholipids sensation by TaPCGR1-3B at different nitrogen supply levels triggered calcium signaling. The different nitrogen supply levels were related to phospholipids degradation and synthesis balance (*16, 20*). Under nitrogen deficient condition, the decrease of PC triggered interaction between TaPCGR1-3B and effector proteins, thus promoting downstream calcium signaling, which was likely to inhibit tillering genes (*Tatab1*) expression. The low expression of *TaPCGR1-3B* in RNAi-positive lines and NIL-*TaPCGR1-3B*^hapI^) made these plants less sensitive to nitrogen deficiency induced phospholipids (such as PC) degradation and Ca^2+^ signaling (Fig. S12), enabled higher tillering gene *Tatab1* expression (Fig. 2E and S10), and ultimately more spike number and grain yield (Fig. 1).

## ACKNOWLEDGMENTS

We thank Prof. Caixia Gao’s laboratory (Institute of Genetics and Developmental Biology, Chinese Academy of Sciences) for developing the transgenic wheat lines.

## Author contributions

Author contributions: Z.H. carried out most of experiments. Y.T. initiated and supervised the project. Z.H. conceived the project. Z.H. and Y.T. designed the experiments. Z.H. performed field experiments. Z.H. and M.H. collected data for traits. Z.H. performed in cellular, molecular and biochemistry experiments. X.H. provided technique support in molecular and biochemistry experiments. X.Z., W.T., M.H., H.L. and J.L. provided technique support in agriculture experiments. Z.H. and Y.Z. analyzed the data and computation. Z.H. wrote the manuscript with input of all other authors. Y.T. revised the manuscript. Competing interests: all authors declare no competing interests. Data and materials availability: all data and material are available on line. Requests for materials should be sent to Y.T.

## List supplementary materials

Materials and methods

Figs. S1 to S12

Tables S1 to S6

## Supplementary materials

### Materials and methods

#### Pant material and growth condition

The winter wheat varieties Xiaoyan 54 (XY54), Jing 411 (J411) and the BC4F5-10 and BC5F1-5 near-isogenic lines of J411//XY54 were planted in the long-term low-nitrogen trial area (2015-2020 growing season) at the Beijing Changping Experimental Base of the Institute of Genetics and Developmental Biology, Chinese Academy of Sciences and the long-term low-nitrogen trial area (2018-2020 growing season) at the Zhao County Modern Agricultural Park Base in Shijiazhuang, Hebei. (2018-2020 growing season). Konong 199 (KN199WT) and various transgenic lines: *TabZIP45-4A* overexpression line, *TaPCGR1*-RNAi lines were grown at the Experimental Station at Dishang, Gao Cheng District, Shijiazhuang, China, of the Institute of Cereal and Oil Crops, Hebei Provincial Academy of Agricultural and Forestry Sciences. The tobacco material used for the transient expression cotransformation experiments was *Nicotiana benthamiana*.

#### Hydrophobic culture condition

The common wheat culture was carried out as previously described with slight modifications (*22*). Seeds were germinated in 0.05 % hydrogen peroxide overnight. The seeds were spread evenly to germinating in moist condition. Seven-day-old seedlings were transferred to plastic pots containing one-liter nutrient solution (0.2 mM KH_2_PO_4_, 1.5 mM KCl, 1.0 mM MgSO_4_.7H_2_O, 1.5 mM CaCl_2_, 1.0 μM H_3_BO_3_, 0.05μM (NH_4_)_6_Mo_7_O_24_.4H_2_O, 0.05μM CuSO_4_.5H_2_O, 1.0 μM ZnSO_4_.7H_2_O, 1.0 Mm MnSO_4_.H_2_O, 0.1 mM FeEDTA(Na), 1.0 mM NH_4_NO_3_). The growth condition was in in 16 hours day - 8 nigh hours cycle at 23°C. The different nitrogen supply levels were 0N (0.0 mM NH_4_NO_3_), 1N (0.5 mM NH_4_NO_3_), 2N (1.0 mM NH_4_NO_3_), 4N (2.0 mM NH_4_NO_3_). All the nutrition solution were refreshed every 4 days and pH was adjusted to 6.0.

#### Plasmid construction

The wheat RNAi transgenic vector: *TaGPCAT* interference line was constructed on *pUBI-RNAi* containing a hairpin structure (kindly donated by Dawen Wang, IGDB). Wheat overexpression transgenic vector: *TabZIP45-4A pro::TabZIP45-4A* was constructed on *163-JIT* backbone (kindly donated by Caixia Gao, IGDB). The above constructed vectors were entrusted to our transformation platform to transform the wheat variety KN199 using the gene gun transformation method as previously described (*22*). Independent transgenic lines were obtained and primers (one end on the vector sequence and the other end on the exogenous sequence) were set at the 5’ and 3’ ends of the respective transgene exogenous sequences, respectively. Positive plants were identified when the size of the sequence fragments amplified at both ends matched that of the plasmid positive control (verified by auxiliary sequencing if necessary) and verified by sequencing. Expression was also identified in field or hydroponic conditions to confirm the success of the respective transgenic events. Primers used were provided in **tableS1**.

#### ChIP-PCR

The procedures were done as precious described (*22, 23*). Briefly, seedling under 0N condition for two weeks and whole seedlings was collected and fixed. The fixed samples were powered and lysis. The supernatant from lysis solution after centrifugated and fragmented by ultrasonic crusher was added with antibody to TabZIP45. The beads were collected after washes and then DNA was isolated. The specific primers were used to evaluated DNA enrichment by quantitative PCR. Primers used were provided in **tableS2**

#### quantitative real-time PCR (qPCR)

Total RNAs were isolated from various tissues according to TRIzol™ Plus RNA Purification Kit (Invitrogen™, USA). The first strand cDNA was synthesized according to cDNA RevertAid First Strand cDNA Synthesis Kit (Thermo Scientific™, USA). The obtained cDNA samples were diluted 20-fold and quantitative PCR experiments were performed according to the instructions of the LightCycler^®^ 480 SYBR Green I Master (Roche Molecular Systems, Inc., CHE). The specificity of the qPCR reaction was confirmed by confirming that the melting curve of the qPCR was a single peak at the end of the reaction. Calculate the mean Ct value. Determine the relative expression of the target gene based on the expression of the corresponding internal reference gene *TaACTIN* (TraesCS1A02G274400). Primers used were provided in **tableS1**

### Subcellular localization

The sequences of the coding regions of *TaPCGR1-3B* gene were cloned into *UBI-163-GFP* (kindly provided by Caixia Gao from IGDB). mCherry-HDEL and PCGR1-GFP were co-transformed into wheat protoplasts. Tobacco protoplasts were isolated after 24 h of injection into Agrobacterium tumefaciens (GV3101) by cloning the sequence of the *TaPCGR1-3B* gene coding region into *PEZR(K)-LC-GFP*. The protoplasts were isolated as previously described (*24*). The protoplasts were observed under a laser confocal microscope (Zeiss LSM710) for fluorescence signals. GFP excitation light range: 500 ∼ 550 nm, RFP/mCherry excitation light range: 565 ∼ 615 nm. For immunoblot staining of ultra-speed separation of different parts of protein was done according to methods previously described (*25*). Proteins were collected and boiled at 95°C for 10 min and spread on PAGE gel. The protein that transformed onto PDVF membrane were labeled with PCGR1 antibody, H^+^-ATPase antibody, TIP antibody, and HSP antibody. Related primers were listed in **table S1**.

### Bimolecular fluorescence complementation (BiFC) assays

CDS sequences of the corresponding genes were amplified by designing adapter primers with attB1 and attB2, and tobacco transient expression vectors were constructed according to the instructions of Gateway^®^Technology (Invitrogen™, USA). The CDS of *TaPCGR1-3B*^HapI^ and *TaPCGR1-3B*^HapII^ was constructed into the vectors for the nitrogen-terminus (N-YFC) of yellow fluorescent protein. *TaPLC, TaPLC-*mutant and *TaGPA* was constructed into the vectors for the nitrogen-terminus (N-YFC) and carbon-terminus (C-YFC) of yellow fluorescent protein, respectively. Sequence verified vector was transformed into *Agrobacterium tumefaciens* GV3101, which was injected into tobacco leaves for 36-48 hours. The leaves were cut and assayed under a laser confocal microscope (Zeiss LSM710) for fluorescence signals. Primers used were provided in **tableS3**

### Electrophoretic mobility shift assay (EMSA)

EMSA was carried out as previously described (*26*). The full length of *TabZIP45-4A* was clone in to the *Pet28a* vector. The recombinant TabZIP45-4A-HIS tag protein was expressed in *E*.*coli* BL21(DE3) strain and purified with Ni-NTA magnetic beads (genscript). The promoter of *TaPCGR1-3B* at 5’UTR region was amplified with biotin tagged primer (Ruibio). The DNA shift in gel was performed according to the LightShift Chemiluminescent EMSA Kit (Thermo Fisher Scientific, Waltham, MA, USA). Related primers were listed in **table S4**.

### Protein purification

*PLC9* and *GPA* were cloned into *pGEX-4T-1* vectors and expressed in *E*.*coli* BL21(DE3) strain. GST Protein were purified with GE sepharose 4B beads (GE, Boston, MA, USA) according to manufacturer’s instructions. Related primers were listed in **table S4**.

### Immunoprecipitation assay (Co-IP) and western blotting

wheat seedlings under 0N condition for two weeks were ground to a fine powder and rapidly dissolved in Radio Immunoprecipitation Assay (*27*)-buffer; 50 ng of GPA and PLC proteins tagged in beads from prokaryotic expression with or without chemicals. 1 hr of mixing at 4°C on a rotary mixer, washed more than 5 times with RIPA-buffer (with or without chemicals). The binding signals was detected by western blotting with GST or PCGR1-3B antibody. Related primers were listed in **table S4**.

### RNA-seq and gene mapping by sequencing

To isolated genes that were responsible for coordination for growth and environmental response, we constructed a BC5F6 population by crossing between a stressed condition sensitive cultivar Jing411(a traditional Chinese winter wheat cultivar) and stresses tolerant cultivar XiaoYan54 (XY54, a widely used cultivar in breeding). The RNA from three paired-pools (one was with more tiller number and the other was less tiller number) were collected. There paired inbreed line from BC4F5-10 and BC5F1-5 (Line27duo-line27shao, line28duo-line28shao, and 2020duo-2020shao) seedlings at tillering stage were collected and RNA was isolated by GeneMarkbio Plant Total RNA Purification Kit (TR02). The cDNA from reverse transcription was sent to library construction. Hiseq-PE150 was used to sequence each library.

The data from sequenced was filter by Trimmomatic (*28*). The resulting sequence was aligned to common wheat genome reference 2.0 by tophat2 (*29*) (--mate-std-dev 50--min-coverage-intron 5 --min-segment-intron 5) and sorted, marked and indexed by samtools (30). SNP were called with samtools/bcftools pipeline (*9*). The SNP were filtered by samtools mpileup (-Bugp -R -t DP,AD,ADF,ADR,SP,INFO/AD,INFO/ADF,INFO/ADR -I -q 50 -Q 14 -s) and bcftools (*31*) and further filter by SWEEP (*32*) when necessary. The SNP depth ratios in different paired pools were calculated according to methods as previously described (*21*). In addition, we confirmed our results by software Triti-Map as previous described (*33*).

### SNPs and splicing variant determination in different haplotypes

Different haplotypes of *TaPCGR1-3B* seedlings were under 0N (0.0 mM NH_4_NO_3_) and 2N (1.0 mM NH_4_NO_3_) conditions for two weeks. The isolated DNA and RNA from these whole seedlings were used to determined SNPs and splicing variant. Related primers were listed in **table S1 and table S5**.

### Transcription activation assay

The promoter of *TaTab1* was cloned into *pBD Gal4 F. GUS* was also constructed into *p163-UBI* vector (kindly given by Caixia Gao). The *pBD-TaTab1-Gal4 F* (firefly luciferase), *p163-UBI-GUS* (internal reference 1) and pTRL-Renilla luciferase (internal reference 2) were co-transferred into wheat protoplasts. GUS intensity assay was performed according to the experimental method previous described (*34*). Luciferase fluorescence values are based on the instructions of the Dual-Luciferase^®^ Reporter Assay System kit (Promega, USA). Related primers were listed in **table S6**.

### Western blotting assay

pull off the comb from 10% NuPAGE Bis-Tris gel (Thermo Scientific™, USA) and put the gel into the electrophoresis instrument, add protein electrophoresis buffer to the electrophoresis tank. Set the voltage at 110 V. Stop electrophoresis once the target protein is expected to have been properly separated according to the protein molecular weight, and remove the gel. The membranes were placed sequentially in the pre-chilled transfer solution as blackboard-sponge-filter paper-gel-PVDF membrane-filter paper-sponge-white board (membranes were soaked in methanol for 5 min before use), and the voltage was set at 200 mA, and the membranes were transferred in an ice bath for 90 min. The PVDF membranes were removed and washed five times with TBST (50 mM Tris-HCl (pH 7.5), 8 g/L NaCl, 0.2 g/L KCl, 0.5% Tween-20) five times for 10 min each. Pour off TBST, add primary antibody in the appropriate proportion in TBST containing 0.5% skimmed milk powder and incubate slowly overnight at 4°C or for 2 h at 37°C. Wash the membrane 3 ∼ 5 times with TBST for 10 min each time. pour off the TBST, add secondary antibody in 0.5% skimmed milk powder in TBST in appropriate proportions and incubate for 1.5 h at 37°C with slow shaking. The membranes were washed 3 ∼ 5 times with TBST for 10 min each time. The membranes were laid flat, ECL reagent was added dropwise to the membranes and photographed with Image Quant LAS 4000.

### Calcium signaling detection

Backcross progeny (BC5F5) sister lines (*TaPCGR1-3B*^HapI^ and control *TaPCGR1-3B*^HapII^) with J411 as the donor parent and XY54 as the rotating parent, *TaPCGR1* reduced expression RNAi+P and the negative line RNAi-N were incubated in greenhouse under high nitrogen conditions (1.0 mM NH_4_NO_3_, HN) for two weeks and then transferred to the test solution, and the Ca^2+^ flow rate in wheat roots (100 μm from the crown) was measured after 10 min of stabilization or incubated under nitrogen-deficient conditions (0.1 mM NH_4_NO_3_, LN) for 5 h and then transferred to the test solution (0.1 mM KCl, 0.1 mM CaCl_2_, 0.1 mM MgCl_2_, 0.5 mM NaCl, 0.3 mM MES, 0.2 mM Na_2_SO_4_, pH 6.0) and then measured the Ca^2+^ flow rate in wheat roots (400 μm from the root crown).

All of the above experiments were performed using a non-invasive technique to determine Ca^2+^ flow rate, and the differences between biological replicates of different strains were counted with maximum Ca^2+^ maximum amplitude after excluding background using an EXCEL sheet. The experiments were converted into the flow rate of ions (pmol cm^-2^ s^-1^) based on the voltage difference. The ion flow rate was calculated according to Fick’s law: J is the ion flow rate (pmol cm^-2^ s^-1^), D_0_ is the diffusion constant, dc is the concentration gradient between two points, and dx is the distance between two points.

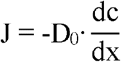

### Calcium staining

Root tip Ca^2+^ staining experiments were performed according to previous described procedures (*35*). Briefly, The Ca^2+^ - sensitive fluorescent dye Fluo-3/AM ester was purchased from Molecular Probes (Eugene, OR, USA). 10-20 mm long wheat root tips were incubated for 2 h at 4°C in staining solution (20 μM Fluo-3/AM ester, 0.2 mM CaCl2 and 50 mM sorbitol) protected from light. Root tips were continued to be incubated for 1-2 h at 20°C in 0.2 mM CaCl_2_ protected from light. Make a chamber of approximately 40 × 15 × 1 mm on the slide to hold the staining solution and root tip, cover with a coverslip, and let stand for 15 min. Observe under an inverted laser confocal microscope (Zeiss LSM710) (excitation light wavelength 488 nm, emission light wavelength 515 nm).

### Immuno-staining

The research steps were operated with reference to methods as previous described (*36*). Briefly, samples were collected in Formaldehyde/alcohol/acetic acid (FAA) solution and vacuum under room temperature. After dehydration, samples were embedded in wax mixture of distearate PEG400 (Sigma Aldrich, USA) and 1-hexadecanol (Sigma-Aldrich). The embedded samples were sectioned into 6 μm slices. The slices were stained with antibody of TaPCGR-3B and secondary antibody Alexa Fluor 488 donkey anti-rabbit IgG. Each slice were mounted on medium with DAPI ProLong Antifade (Life Technologies). Images of each slice were taken under an inverted laser confocal microscope (Zeiss LSM710) (excitation light wavelength 488 nm, emission light wavelength 515 nm).

### Evolutionary tree building

The homologous protein sequences of TaPCGR1 were downloaded from the database (http://plants.ensembl.org and https://www.rcsb.org). Evolutionary distances were calculated and evolutionary trees were constructed using MEGA 6.0 software for neighbor-joining.

### Statistical analysis

All data were counted and plotted using GraphPad Prism 8 or Microsoft office 2016. p-values less than 0.05 were considered to be significantly different. Appropriate statistics were selected according to the significance of the orthogonal distribution of the data and the conditions of processing.

**Fig.S1.**
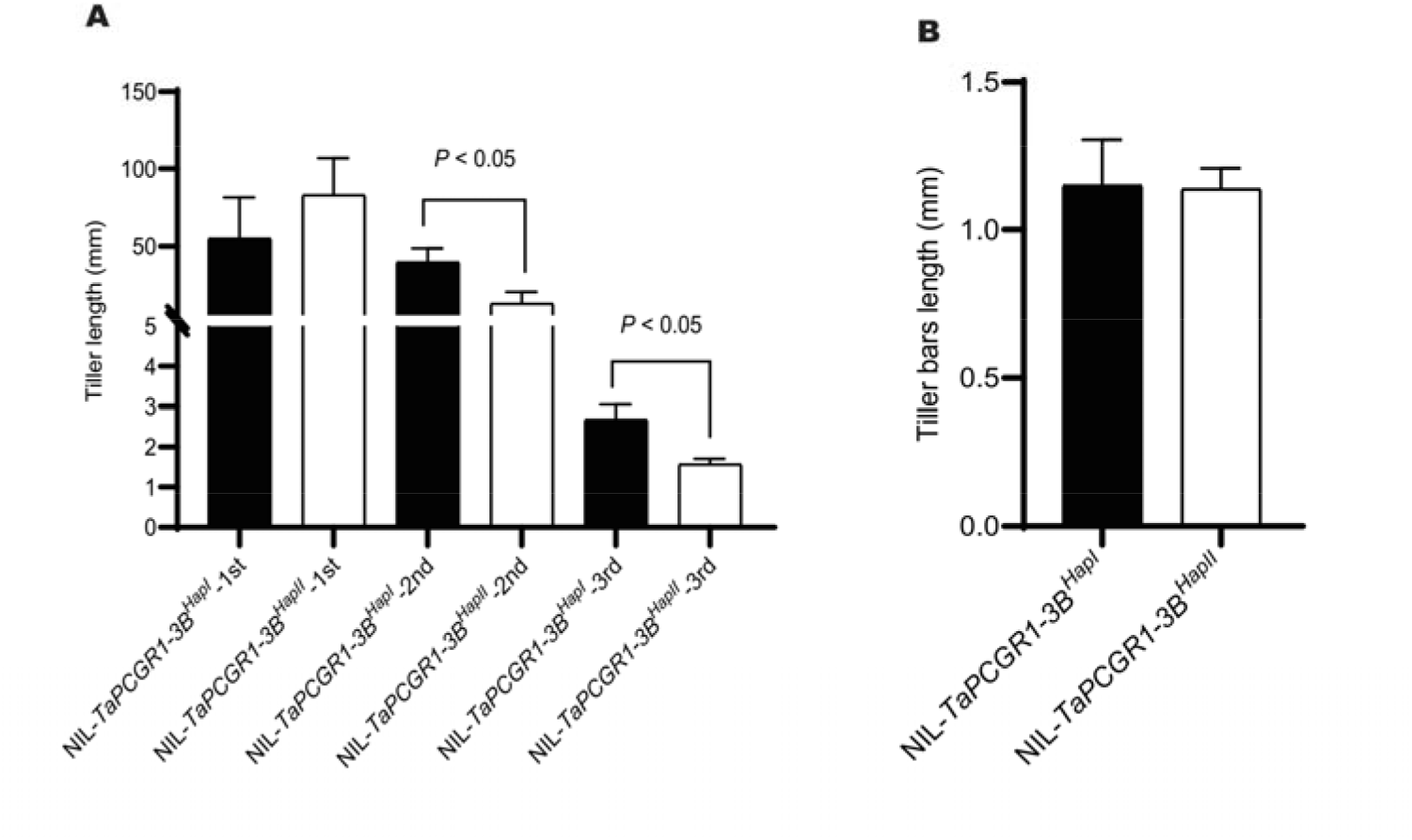
*TaPCGR1-3B*^HapI^ promoted tiller elongation. (**A**) Tiller buds length in HN. (**B**) Tiller buds number in LN. Seedlings were grown under high N conditions (1.0 mM NH4NO3, HN) or (0.1 mM NH4NO3, LN) for two weeks and samples were collected. Data are mean ± S. E. (n = 8) and *P* values in E-H were from paired Student’s *t*-test. 1st is first tiller, 2nd is second tiller. Isogenic lines *TaPCGR1-3B*^HapI^ and *TaPCGR1-3B*^HapII^ were generated by introducing *TaPCGR1-3B*^HapI^ from J411 into XY54 (BC5F5)

**Fig.S2.**
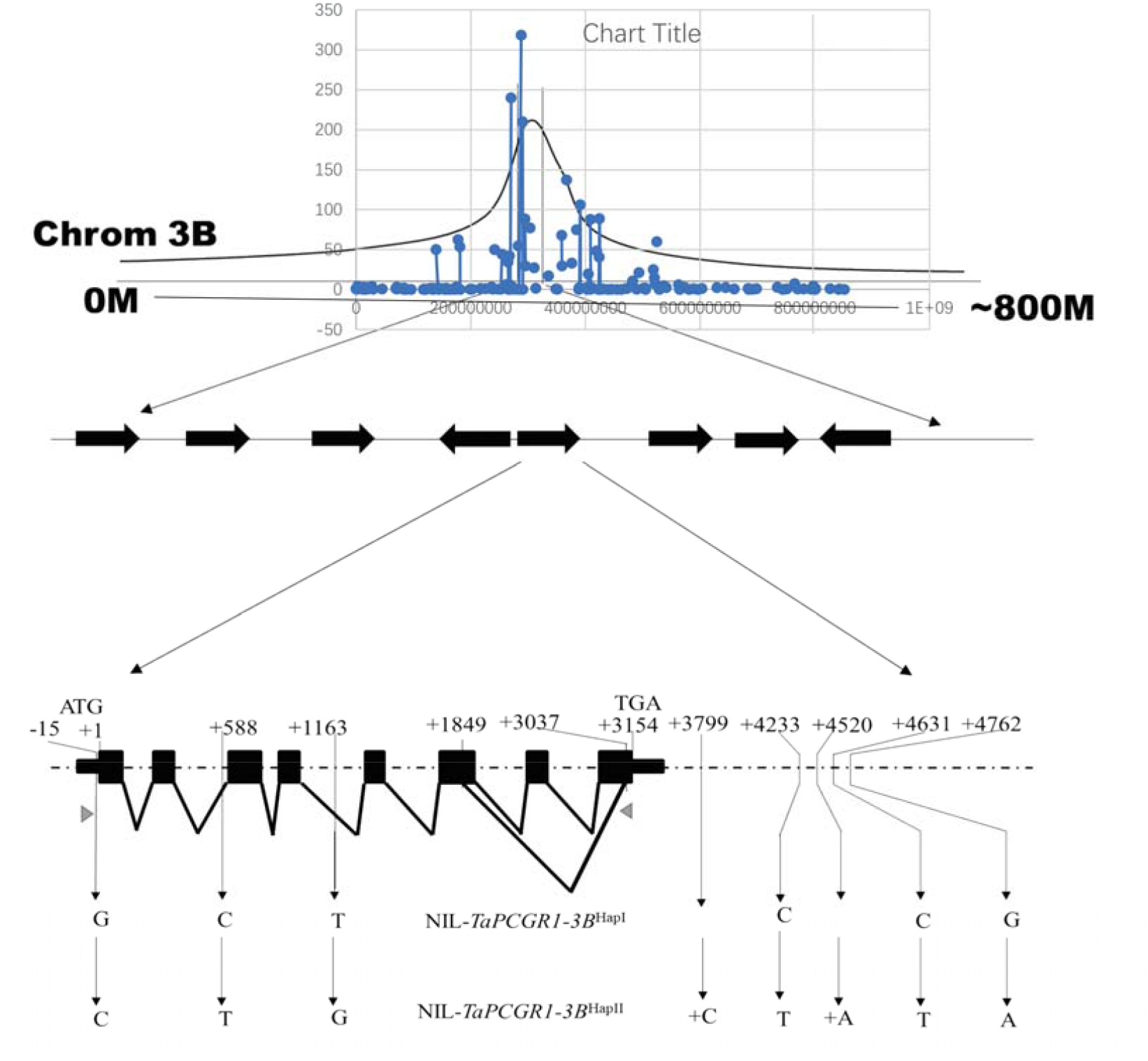
Genetic mapping for membrane protein *TaPCGR1-3B*. The isolation of *TaPCGR1-3B* by mapping of sequencing. the mutation sites indicated by positions (numbers) and SNP (with arrow headed lines and character). Black rectangles, exons; thin rectangles, UTRs; concave lines, splicing joining between exons; gray triangles, RT-PCR primers positions.

**Fig.S3.**
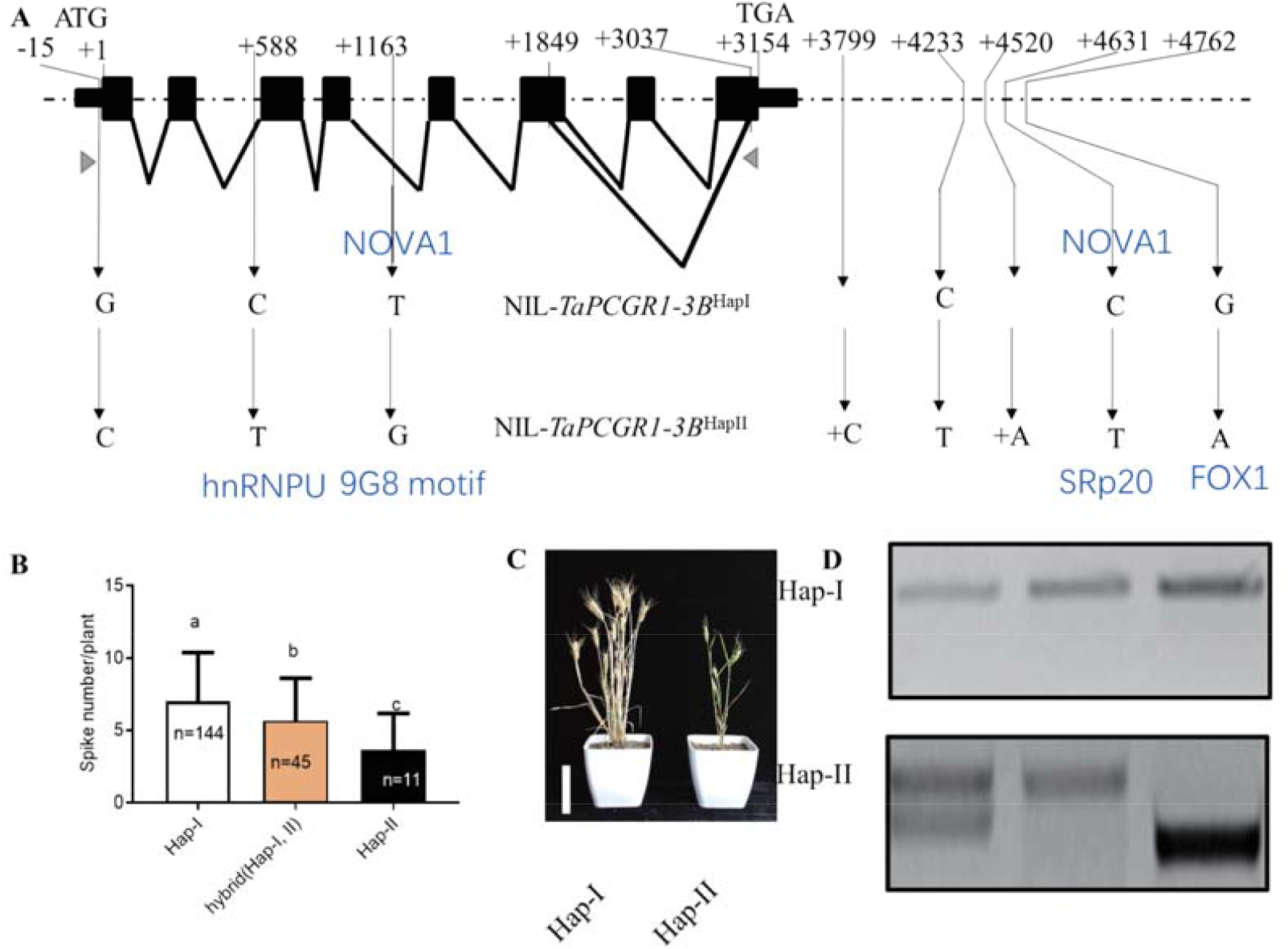
*TaPCGR1-3B* haplotype trait and splicing variants. (**A**) Two haplotypes of *TaPCGR1-3B* in separating near isogenic lines. The isogenic lines NIL-*TaPCGR1-3B*^HapI^ (conferring NIL-*TaPCGR1-3B*^HapI^ without splicing variant) and NIL-*TaPCGR1-3B*^HapII^ (conferring NIL-*TaPCGR1-3B*^HapII^ with splicing variant) were screened from J411//XY54 backcrosses (BC5F5-7). Black rectangles, exons; thin rectangles, UTRs; concave lines, splicing joining between exons; gray triangles, RT-PCR primers positions. Blue letters, splicing binding factors change in each SNP respectively (**B**) The corresponding spike number per plant of two isogenic lines in 2018-2019 growing season in Beijing. (**C**) Representative plants selected from more than ten replicates. Scale bar, 15 cm. (**D**) Two splicing variants of two haplotypes.

**Fig.S4.**
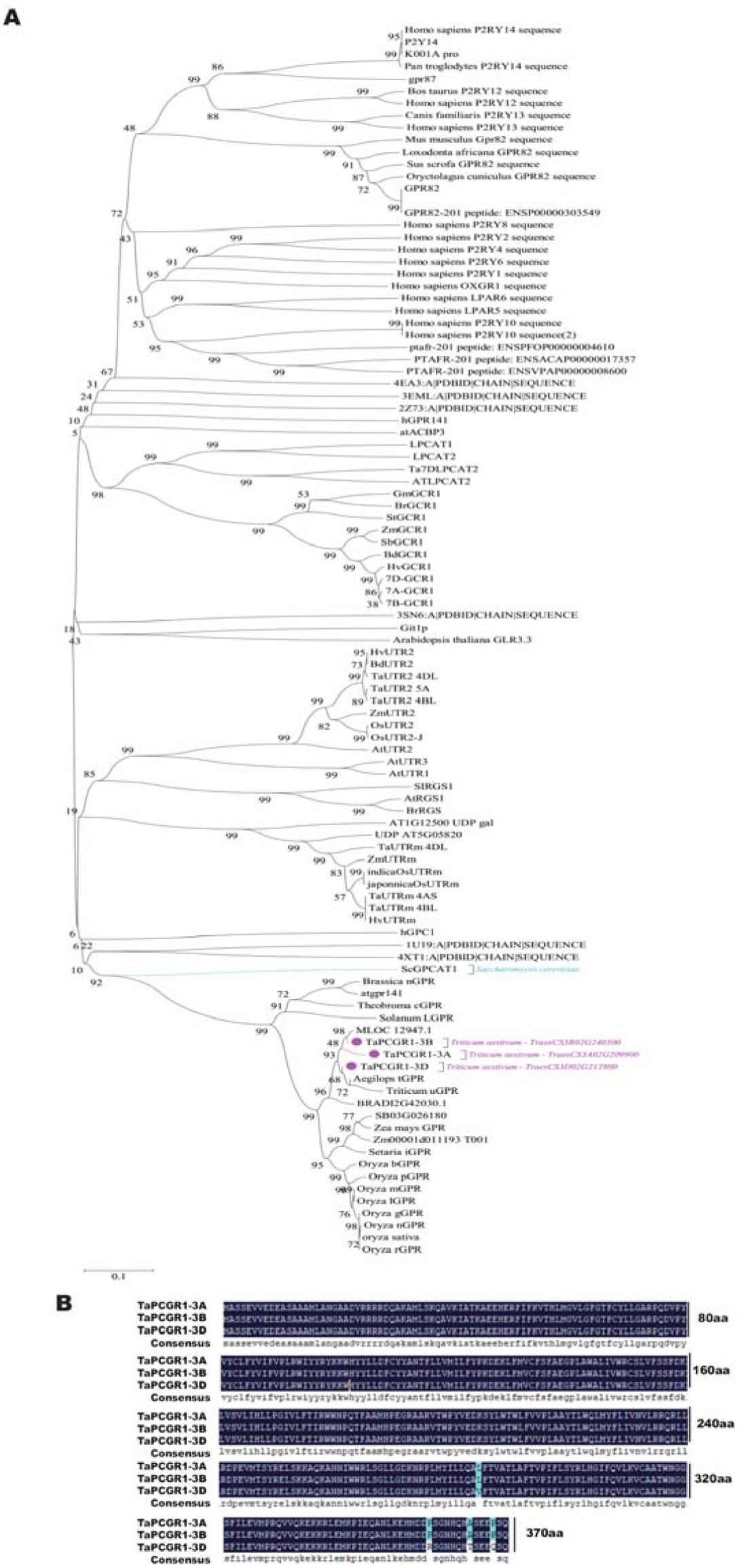
PCGR1 family and TaPCGR1 sequence. **(A)** Phylogeny tree of PCGR1 protein family. The Phylogeny tree is illustrated by MEGA 6.0 by neighbor-joining method. (**B**) Coding sequences of *TaPCGR1-3A, -3B* and -*3D* cloned in KN199, which was generated by DNAMAN (LynnonBiosoft, USA).

**Fig.S5.**
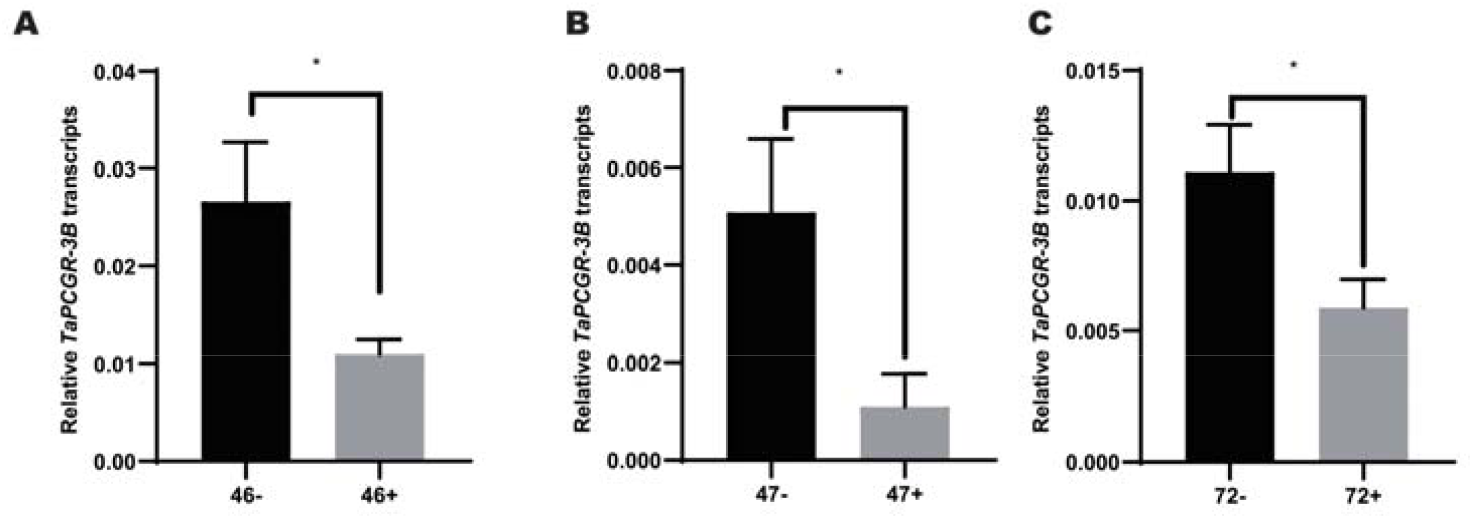
Three independent *TaPCGR1* transgenic lines. The abundance of *TaPCGR1-3B* transcripts in leaves of transgenic positive lines (46+, 47+ and 72+) and negative control lines (46-, 47- and 72-) at tillering stage under nitrogen deficient conditions (30 kg N/ha). The genes expression level was normalized to that of *TaACTIN* as internal control. Data are mean ± S. E. (n = 8).

**Fig.S6.**
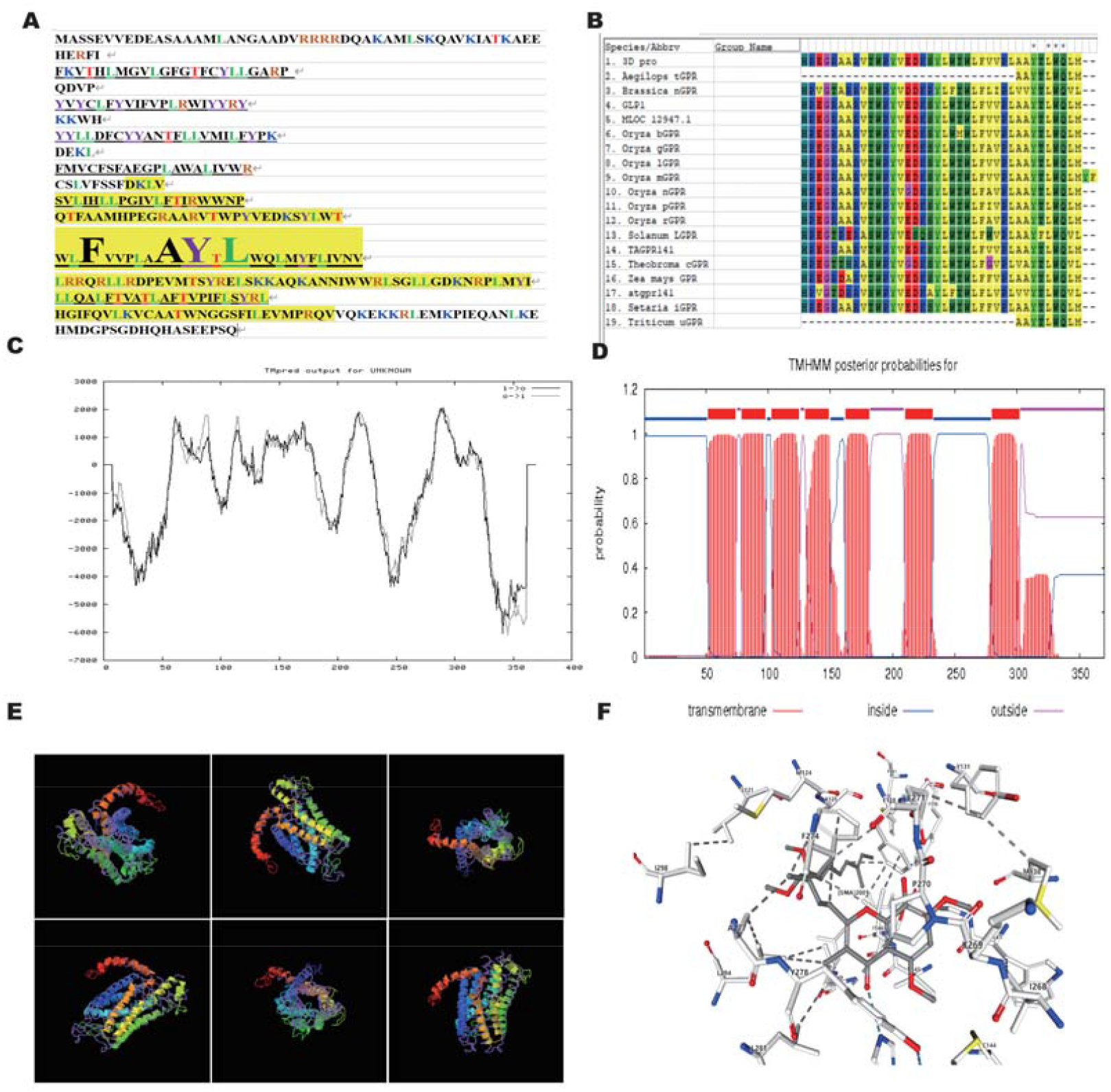
TaPCGR1-3B structure prediction. **(A)** The yellow region indicates the deletion of aa in the short splicing variant. (**B**) The conservation aa sites of TaPCGR1-3B in diverse species. (**C**) prediction was produced byTMpred (https://embnet.vital-it.ch/software/TMPRED_form.html) (**D**) prediction was produced by TMHMM (http://www.cbs.dtu.dk/services/TMHMM). (**E**) Structure prediction of wild type TaPCGR1-3B protein (predicted by I-TASSER). (**F**) The interaction sites between protein and chemical by thread alignment of machine learning [predicted by FALCON (http://protein.ict.ac.cn/TreeThreader/) and structure of binding sites from Protein Data Base].

**Fig.S7.**
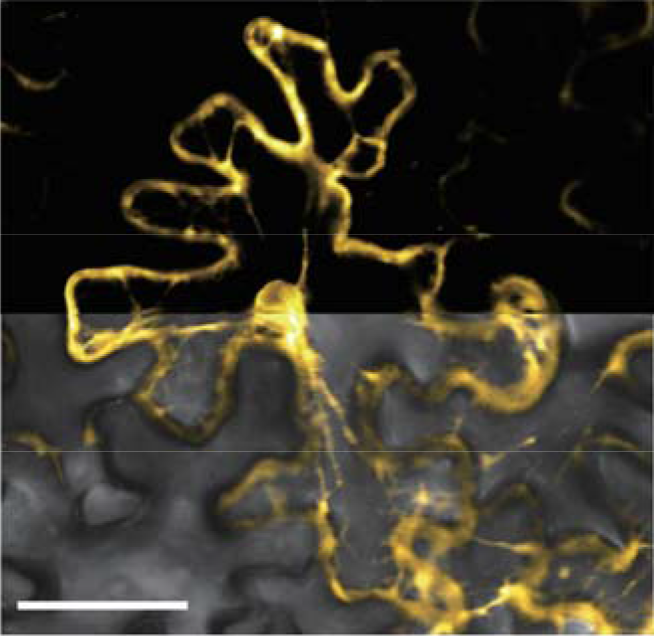
TaPCGR1-3B mutant (short variant of *TaPCGR1-3B*^HapII^) was targeted by TaGPA1. BiFC experiment: gene1-YFP C terminal part (TaGPA1**-**Y^C^) and targeting gene2-YFP N terminal part (TaPCGR1-3B mutant **-**Y^N^) co-expressed in tobacco (*Nicotiana tabacum*) leaves and then was scanned with confocal microscope. Scale bar, 100 μm.

**Fig.S8.**
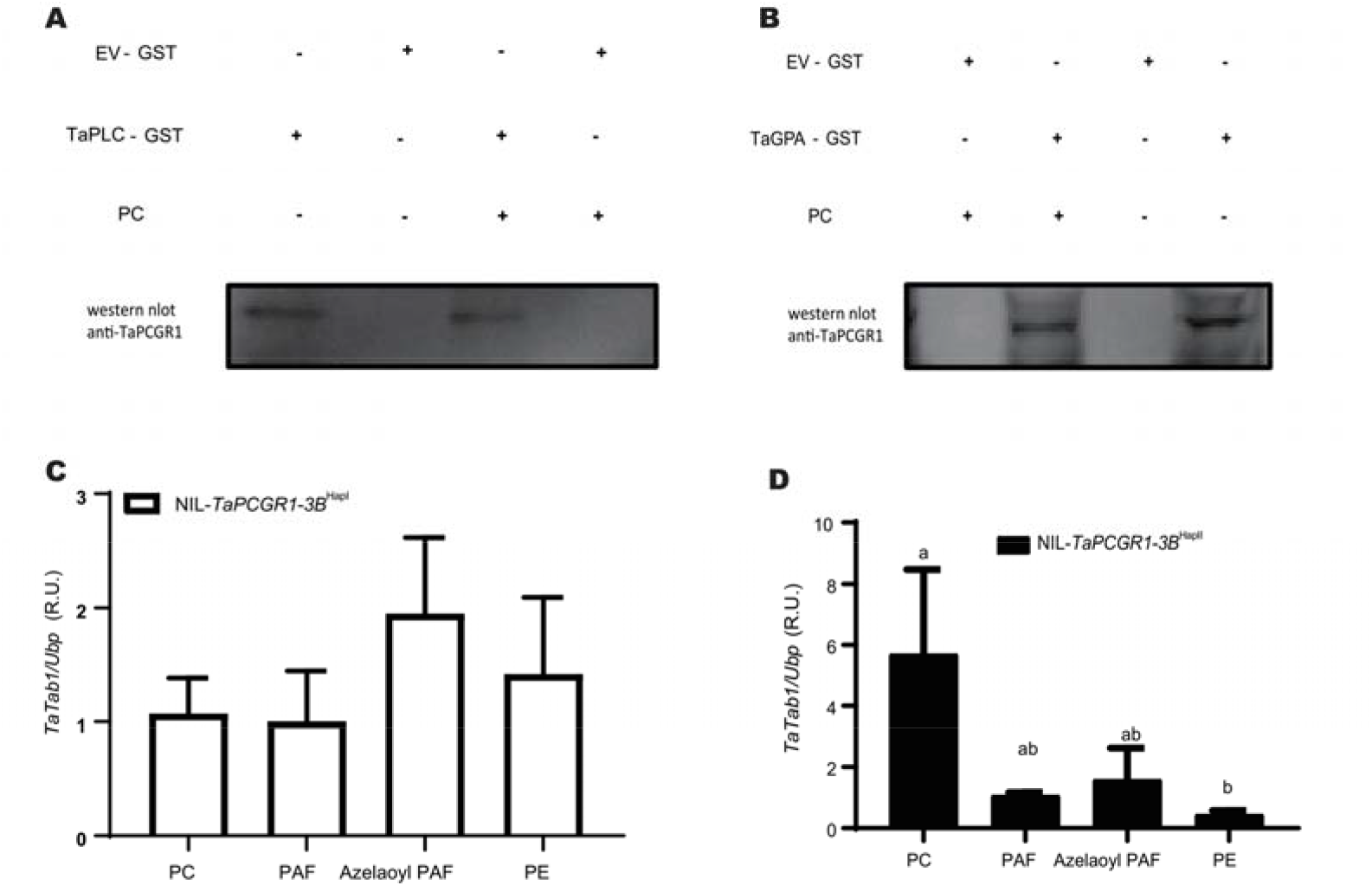
PCGR1 interaction with effectors proteins was affected by PC. (**A)** Co-immunoprecipitation assay: Purified GST-TaGPA1 or GST-EV was incubated with KN199WT root tissue lysate with or without PC, and then was detected by western-blot. (**B**) Co-immunoprecipitation assay: Purified GST-TaPLC9 or GST-EV was incubated with KN199WT root tissue lysate with or without PC, and then was detected by western-blot. (**C**-**D**) Relative *TaTab1*-luciferase fluorescence (R.U., relative units). Protoplasts were isolated from NIL-*TaPCGR1-3B*^HapI^ (C) and NIL-*TaPCGR1-3B*^HapII^ (D). Isogenic lines NIL-*TaPCGR1-3B*^HapI^ and NIL-*TaPCGR1-3B*^HapII^ are generated by introducing *TaPCGR1-3B*^HapII^ from J411 into XY54 (BC5F6). Fluorescence was normalized first with internal control (Ubq-GUS or Renilla luciferase). The value was further normalized to negative control value respectively. Data are mean ± S. E. (n = 4) and different letters denote statistically significant differences (*P* < 0.05) from one-way ANOVA multiple comparisons.

**Fig.S9.**
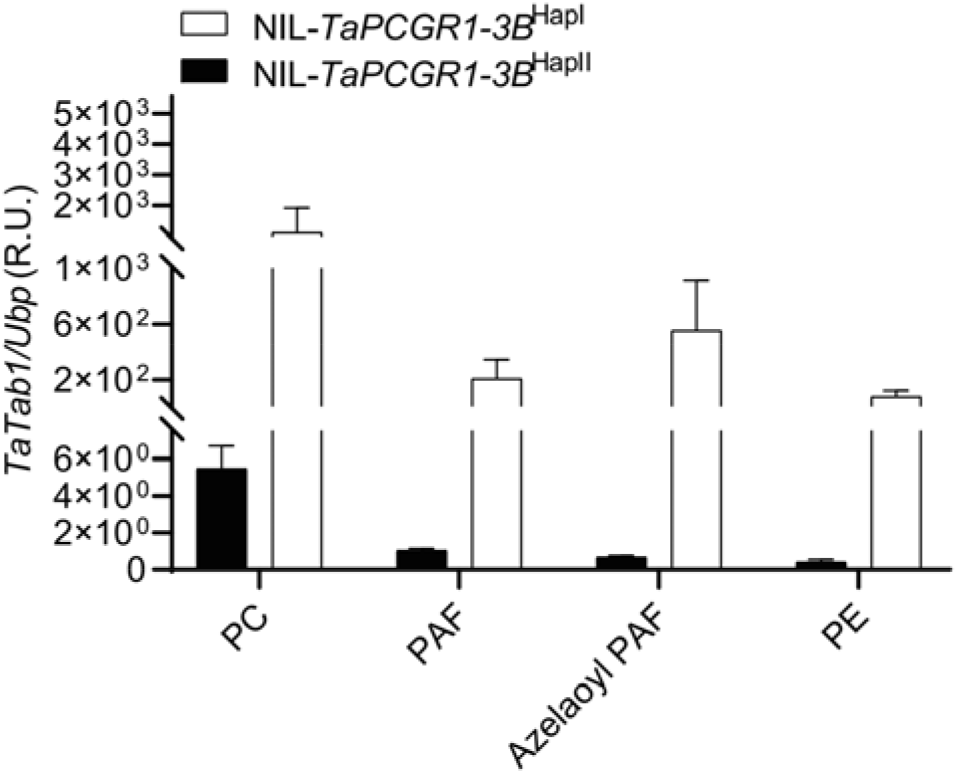
*Tatab1* was differently regulated by phospholipids in different haplotypes of *TaPCGR1*. Relative *TaTab1*-luciferase fluorescence (R.U., relative units). Protoplasts were isolated from NIL-*TaPCGR1-3B*^HapI^ and NIL-*TaPCGR1-3B*^HapII^. Isogenic lines NIL-*TaPCGR1-3B*^HapI^ and NIL-*TaPCGR1-3B*^HapII^ are generated by introducing *TaPCGR1-3B*^HapII^ from J411 into XY54 (BC5F6). Fluorescence was normalized first with internal control (Ubq-GUS or Renilla luciferase). The value was further normalized to negative control value respectively (The fluorescence intensity of NIL-*TaPCGR1*-*3B*^hapI^ by solvent stimulation was set as 1). Data are mean ± S. E. (n = 4) and different letters denote statistically significant differences (*P* < 0.05) from one-way ANOVA multiple comparisons.

**Fig.S10.**
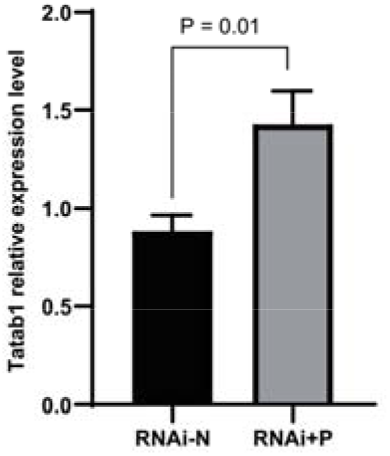
*TaTab1* was acted downstream of *TaPCGR1*. The abundance of *Tatab1* transcripts in leaves of transgenic positive lines and negative control lines at tillering stage under nitrogen deficient conditions (30 kg N/ha). The genes expression level was normalized to that of *TaACTIN* as internal control. Data are mean ± S. E. (n = 8).

**Fig.S11.**
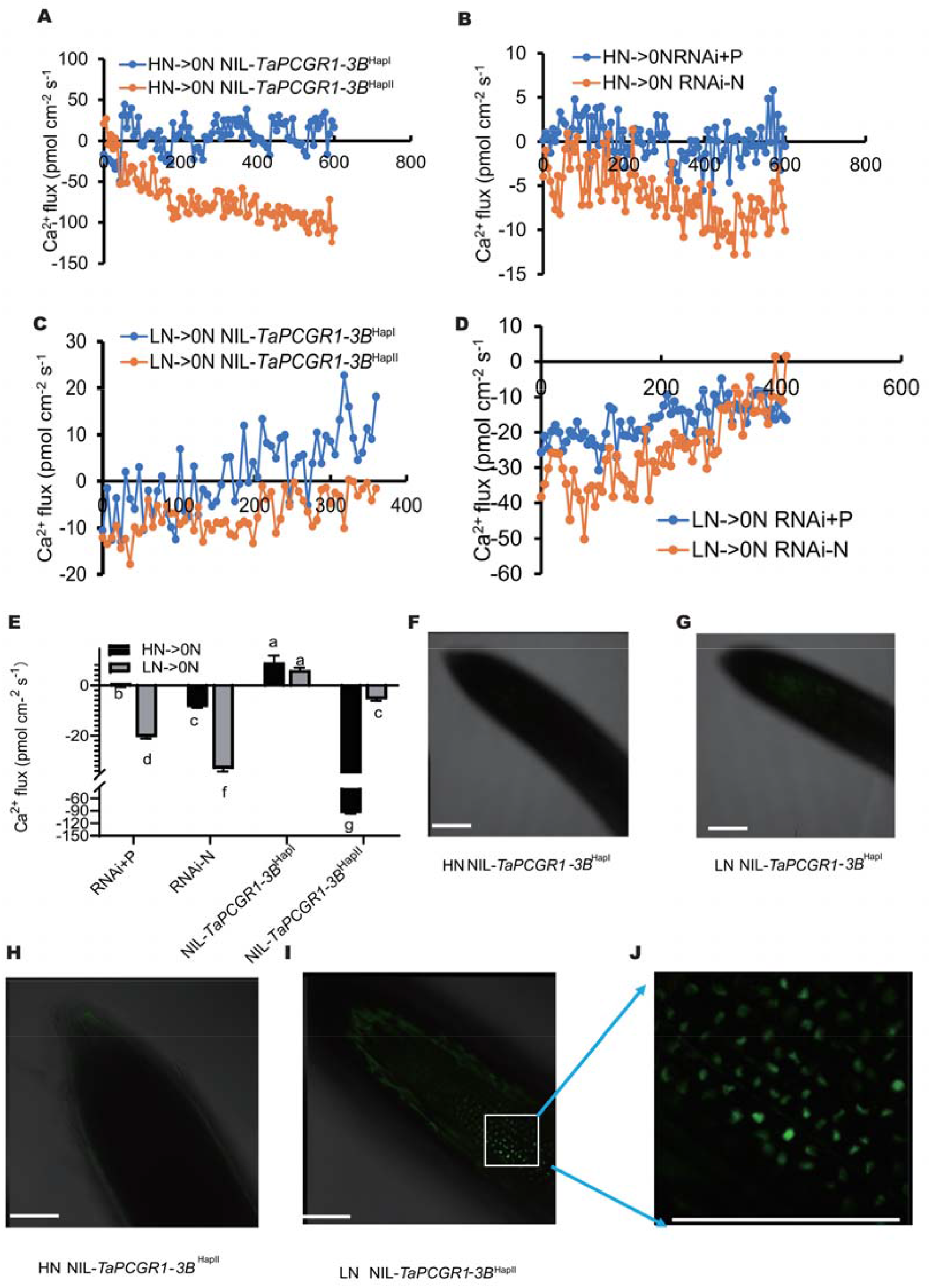
*TaPCGR1* mediated nitrogen level sensation. (**A-B**) Wheat seedlings were grown under high N conditions (1.0 mM NH_4_NO_3_, HN) for two weeks, then were transferred into the test buffer (0.1 mM KCl, 0.1 mM CaCl_2_, 0.1 mM MgCl_2_, 0.5 mM NaCl, 0.3 mM MES, 0.2 mM Na_2_SO_4_, pH 6.0) for 10 min before Ca^2+^ flux measurement at 100 μm from root tip. (**C-D**) Wheat seedlings under HN conditions two weeks were transferred into a low N solution (0.1 mM NH_4_NO_3_, LN) for another five hours before test at 400 μm from root tips. (A and C) NIL-*TaPCGR1-3B*^HapI^ and NIL-*TaPCGR1-3B*^HapII^ generated by introducing *TaPCGR1-3B*^HapII^ from J411 into XY54 (BC5F6). (B and D) RNAi positive line (RNAi+P) and RNAi negative line (RNAi-N). (**E**) statistical analysis in (A-D). (**F-G**) Ca^2+^ staining in the roots of NIL-*TaPCGR1-3B*^HapI^ wheat seedlings under HN (F) conditions or LN (G) conditions. (**H-J**) Ca^2+^ staining in the root of NIL-*TaPCGR1-3B*^HapII^ wheat seedlings under HN (H) conditions or LN (I-J) conditions. Wheat seedlings (G, I and J) were under HN conditions for two weeks before transferred to the low N solution (0.1 mM NH_4_NO_3_, LN) for another 14 hours. Scale bar, 100 μm.

**Fig.S12.**
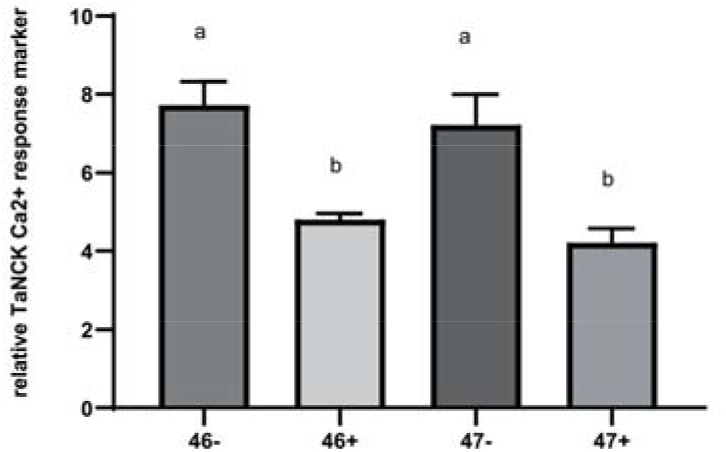
Calcium responsive gene *TaNCK1* was acted downstream of *TaPCGR1*. The abundance of *TaNCK1* transcripts in leaves of transgenic positive lines and negative control lines at tillering stage under nitrogen deficient conditions (30 kg N/ha). The genes expression level was normalized to that of *TaACTIN* as internal control. Data are mean ± S. E. (n = 8)

